# IGF-1 therapy improves muscle size and function in experimental peripheral arterial disease

**DOI:** 10.1101/2022.07.31.502209

**Authors:** Gengfu Dong, Chatick Moparthy, Trace Thome, Kyoungrae Kim, Terence E. Ryan

## Abstract

Lower extremity peripheral arterial disease (PAD) has continued to increase in prevalence over the past several decades, yet therapeutic development has remained stagnant. Skeletal muscle health and function has been strongly linked to quality of life and medical outcomes in PAD patients. Using a rodent model of PAD, this study demonstrates that treatment of the ischemic limb with adeno-associated virus-mediated expression of insulin-like growth factor 1 (IGF1) significantly increases muscle size and strength, without improving limb hemodynamics. Interestingly, the effect size of IGF1 therapy was larger in female mice compared to their male counterparts, where substantial improvements in muscle specific force and a reduction in the progression of limb necrosis were observed. These findings indicate that clinical trials should carefully examine sex-dependent effects in experimental PAD therapies.

## INTRODUCTION

Peripheral arterial disease (PAD), caused by atherosclerotic narrowing or occlusion of blood vessels in the lower extremities (1), affects an estimated 200 million people throughout the world (2,3) making it the third-leading cause of the cardiovascular death (3). A small proportion of PAD patients develop chronic limb threatening ischemia (CLTI), the condition’s most severe manifestation, which is characterized by chronic ischemic rest pain that can occur with or without non-healing wounds, ulcers, or gangrene (4). Based on the symptom severity, CLTI patients have an increased risk of amputation of the lower extremity and cardiovascular mortality compared to PAD patients without CLTI (4,5). Primary treatments for CLTI include endovascular and revascularization surgeries aimed to improve blood flow to the affected limb (6,7). Despite advances in surgical techniques, the complexity of CLTI pathobiology and the moderate-to-high risk classification of most CLTI patients contributes to unfavorable failure rates (8-11). Unfortunately, non-surgical medical therapy is limited risk factor-modifying treatments such as anti-thrombotics, anti-hypertensives, lipid-lowering, and glucose-controlling medications. These therapies are effective at reducing cardiovascular events such as myocardial infarction and stroke, but unfortunately do not significantly improve walking performance or functional ability in PAD patients. Thus, there is a substantial need to develop new medical therapies that can be effective at improving limb function and outcomes in PAD/CLTI.

Gene and cell-based regenerative therapies have emerged as leading technologies for the treatment of numerous human diseases. Specifically, adeno-associated viruses (AAV), a non-pathogenic DNA virus, have been successfully used to treat spinal muscular atrophy and retinal dystrophy in humans, while numerous other AAV-based gene therapies are currently in clinical trials. Several clinical trials have tested gene or cell therapies in PAD/CLTI, however, to date no trials have proven effective (12-17). The lack of success in translating gene and cell-based therapies to CLTI patients is multifactorial and undoubtedly involves the heterogeneity of patient characteristics. Further to this, early-stage preclinical models may not have adequately modeled the CLTI condition. The majority of gene and cell-based therapies have been applied in mouse and rat models of hindlimb ischemia (femoral artery ligation), however the therapeutic has been most commonly applied either prior to, or immediately after surgery before the severe ischemic tissue pathology has developed (18-27). Obviously, the timing of these treatments is not directly relevant to the PAD/CLTI patient who is generally referred to a specialist following the development of symptoms. Moreover, most studies have utilized the C57BL6 genetic background which is known to recover rapidly from hindlimb ischemia and does not develop the CLTI phenotype (24,25,28-32) and is also resistant to acute ischemic injury when compared to the BALB/c mouse (33).

In this study, we first aimed to optimize the dosing and injection strategy for the delivery of self-complimentary AAV9-based gene therapy vectors to the murine CLTI limb. We hypothesized that strategies with lower injection volumes and higher viral titers would produce the greatest levels of viral infection with minimal side effects stemming from the injection. Next, we employed the most effective AAV9 treatment strategy for CLTI to examine the potential therapeutic effects of insulin-like growth factor 1 (IGF1), which is known to promote muscle hypertrophy and enhance regeneration from injury in non-CLTI conditions (34-36). We hypothesized that PAD/CLTI mice treated with scAAV9-IGF1 would exhibit improved limb recovery compared to scAAV9-GFP controls.

## MATERIALS AND METHODS

### Animals

Experiments were conducted on 8 to10-week-old BALB/cJ male and female mice (n=56) purchased from Jackson Laboratories (strain #: 000651). All mice were housed in a temperature- (22°C) and light-controlled (12h light/12h dark) room and maintained on standard chow diet (Envigo Teklad Global 18% Protein Rodent Diet 2918 irradiated pellet) with free access to food and water. All animal experiments adhered to the Guide for the Care and Use of Laboratory Animals from the Institute for Laboratory Animal Research, National Research Council, Washington, D.C., National Academy Press. All procedures were approved by the Institutional Animal Care and Use Committee of the University of Florida (Protocol 202010121).

### Animal model of PAD/CLTI

Femoral artery ligation (FAL) (37) was performed by anesthetizing mice with intraperitoneal injection of ketamine (90 mg/kg) and xylazine (10 mg/kg) and surgically inducing unilateral hindlimb ischemia by placing ligations on the femoral artery just distal the inguinal ligament and immediately proximal to the saphenous and popliteal branches. Because most female patients with PAD/CLTI are older in age and post-menopausal, female mice underwent bi-lateral oophorectomy seven to ten days prior to FAL. Buprenorphine (0.05 mg/kg) was given post-operatively for analgesia. Limb necrosis was scored using the following scale: grade 0, no necrosis in ischemic limb; grade I, necrosis limited to toes; grade II, necrosis extending to dorsum pedis; grade III, necrosis extending to crus.

### AAV Plasmid Construction

All experiments herein were performed using self-complimentary (sc) AAV vectors. The scAAV-CMV-GFP plasmid was purchased from Cell Biolabs (Cat. No. AAV410). The insulin-like growth factor 1 (IGF1) coding sequence was PCR amplified (CloneAmp, Takara Bio; Cat. No. 639298) from cDNA generated from a rat gastrocnemius muscle. The scAAV-CMV-GFP plasmid was digested with EcoRI and HpaI (New England Biolabs; Cat. Nos. R3101 and R0105) and the linearized scAAV-CMV plasmid without the GFP was gel purified. The rat IGF1 coding sequence was then inserted into the linearized scAAV-CMV plasmid using In-Fusion cloning (Takara Bio; Cat. No. 639650). The resulting plasmids for scAAV-CMV-GFP and scAAV-CMV-IGF1 were packaged using AAV2/9 serotype by Vector Biolabs (Malvern, PA).

### AAV Delivery to the CLI Limb

scAAV9 was delivered via intramuscular injections of the hind limb gastrocnemius, tibialis anterior (TA), extensor digitorum longus (EDL), and flexor digitorum brevis (FDB) muscles seven days after FAL surgery. To determine the optimal AAV9 dosage and volume for the CLTI limb, four injection strategies were compared to a sham injection (insertion of needle without injection) as shown in Figure 1A. Mice were randomized to treatment groups at the time of injection based on limb necrosis scores. Experimenters responsible for performing all outcomes measures were blinded to the treatment group of the mice. Female animals received the same virus dosage, but the injection volume was decreased based on the percent difference in muscle weights compared to males as follows: TA: 25μL, EDL: 12μL gastrocnemius: 46μL FDB: 8μL.

**Figure 1.**
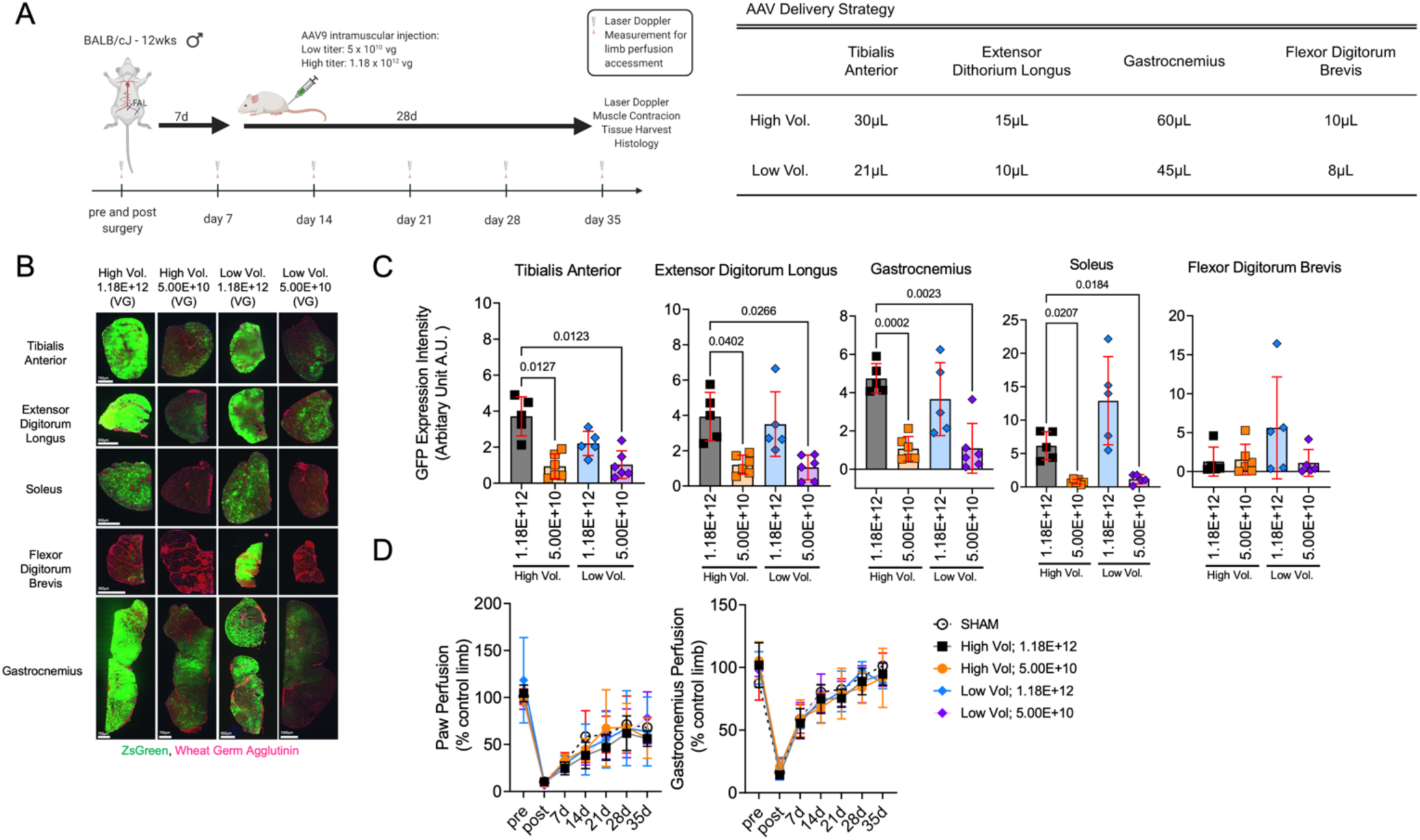
Establishment of the optimal AAV dosing and delivery strategy for the critically ischemic limb. (A) Graphical abstract of study design created using Biorender.com and AAV injection volumes. (B) Representative images of tibialis anterior, extensor digitorum longus, soleus, flexor digitorum brevis, and gastrocnemius muscles injected with scAAV9-CMV-GFP (n=5-6/group) 35 days post-FAL surgery. (C) Quantification of the GFP expression intensity (integral of area and intensity) across ischemic hindlimb muscles (n=5-6/group). (D) Laser Doppler paw and gastrocnemius muscle perfusion recovery (n=3-6/group). Error bars represent SD. Panels C and D were analyzed using Brown-Forsythe and Welch ANOVA with Dunnett’s T3 multiple comparisons testing.

### Assessment of Limb Perfusion with Laser Doppler

Limb perfusion was measured using a laser Doppler flowmeter (moorVMS-LDF; Moor Instruments, Wilmington, Delaware) before and immediately following FAL surgery, as well as weekly measurements until 35 days after the FAL surgery. Under ketamine/xylazine anesthesia, the mice were placed on a water-circulating pad at 37°C, and the leg hair was shaved using a pen trimmer. Once the mouse’s breathing pattern was stable, the laser Doppler probe was placed against the skin of the plantar skin proximal to the footpad, mid-belly of the TA muscle, and the mid-belly of the lateral side of the gastrocnemius muscle to access the peripheral tissue blood flow. The data were collected for 30 seconds, and the average perfusion flux values were calculated. Limb perfusion was presented as a percentage of the non-ischemic contralateral limb.

### Assessment of Hindlimb Grip Strength

Unilateral hindlimb grip strength was measured using a Grip Strength Test Instrument (BIOSEB; Model No. BIO-GS3) prior to FAL surgery and 35 days post-surgery. The mice were allowed to firmly grip the metal T bar and the mice were pulled backward horizontally with increasing force until release the T bar. Three trials were performed per limb with 30 seconds of rest between each trial, and the highest force was recorded.

### Nerve-mediated Muscle Contractile Function

The maximal twitch and tetanic force levels of the tibialis anterior (TA) muscle were measured *in-situ* using stimulation of the sciatic nerve as previously described (38). Briefly, the TA tendon was tied using a 4-0 silk suture attached to the lever arm of the force transducer (Cambridge Technology; Model No. 2250). Muscle contractions were elicited by stimulating the sciatic nerve via bipolar electrodes using square wave pulses of 0.02ms (701A stimulator; Aurora Scientific). Data collection and servomotor control were manipulated using a Lab-View-based DMC program (version v5.500). TA muscles were stretched to the optimal length first. Next, three isometric tetanic forces were acquired using a train of 150Hz, 500ms supramaximal electrical pulses at the optimal length in the muscle, and the highest force among the three measurements was recorded as the peak tetanic force. One minute of rest was provided between each tetanic contraction. Following tetanic contractions, three isometric twitch contractions (1Hz) were performed, and the highest force was recorded as the peak twitch force.

### Skeletal Muscle Histology and Immunofluorescence Micoscopy

Skeletal muscle morphology was assessed by standard light microscopy with hematoxylin and eosin (H&E) staining as previously described (39). Skeletal myofiber cross-sectional area (CSA) and perfused capillary density were assessed by immunofluorescence microscopy as previously described (39). In brief, 10 to 12-µm-thick transverse sections were cut from TA, EDL, gastrocnemius, soleus and FDB muscles. Muscle sections were fixed with 4% paraformaldehyde (in PBS) for five minutes at room temperature followed by 30 minutes of incubation with Alexa647-conjugated Wheat Germ Agglutin (ThermoFisher Scientific; Cat. No. W32466, 1:200 dilution) or overnight incubation at 4°C with a primary antibody for laminin (Millipore-Sigma; Cat. No. L9393, 1:100 dilution) following with Alexa-Fluor secondary antibody incubation (ThermoFisher Scientific; Cat. No. A21428, 1:250 dilution). Images were obtained at 20x magnification using an Evos FL2 Auto (ThermoFisher Scientific) and tiled images of the entire muscle were used for analysis. 50-µL of Griffonia simplicifolia I isolectin (GSL) B4 (Vector laboratories; Cat. No. DL1207) was given to mice through a retro-orbital injection and allowed to circulate systemically for 90 minutes to label perfused capillaries. All images were coded, and analysis was performed by blinded investigators. The level of GFP infection was quantified using Image J as the integral of the area and fluorescence intensity obtained by manually circling the entire muscle. Myofiber CSA were quantified using the Muscle J program on laminin-stained muscle sections (40). Using Image J, number of perfused capillaries were measured by manual counting of isolectin-positive vessels within the entire muscle section. Similarly, the non-fiber area of the skeletal muscle was quantified by manually adjusting the threshold of the H&E staining images to identify and measure the area of non-myofiber tissue within the muscle section. The ischemic lesion area was quantified by manually tracing areas within the muscle section that showed clear signs of ischemic injury (i.e., necrotic fibers, centralized nuclei, infiltrating mononuclear cells) using H&E images in Image J. The percentage of myofibers with centralized nuclei were quantified by manually counting of fibers in six randomly selected 20x images per muscle section for the non-surgical limb, and the entire muscle section in the surgical limb.

### RNA-isolation and qRT-PCR

Total RNA was extracted from gastrocnemius muscle using TRIzol (Invitrogen; Cat. No.15596018). The muscle sample was homogenized with TRIzol using PowerLyzer 24 (Qiagen), and RNA was isolated using Direct-zol RNA MiniPrep kit (Zymo Research; Cat. No. R2052) following the manufacturer’s direction. RNA quantity and quality was assessed as previously described (41). cDNA was generated from 200ng of RNA using PrimeScript RT reagent Kit with gDNA Eraser (Takara Bio; Cat. No. RR047A) according to the manufacturer’s directions. Real-time PCR (RT-PCR) was performed on a Quantstudio 3 (ThermoFisher Scientific) using GoTaq qPCR Master Mix (Promega; Cat. No. A6001) and SYBR green primers for rat-IGF (F:5’-CTACAAAGTCAGCTCGTTCCA-3’, R: 5’-TCTTGTTTCCTGCACTTCCTC-3’, Integrated DNA Technologies). β-actin (F:5’-GGCTGTATTCCCCTCCATCG-3’, R: 5’-CCAGTTGGTAACAATGCCATGT-3’, Integrated DNA Technologies) was used as the housekeeping gene. Relative gene expression was calculated using 2^- ΔΔCT^ from the GFP control group.

### Statistical analysis

Data are presented as mean ± SD. Normality of data was tested with the Shapiro-Wilk test. Because of the sample size in experiments comparing AAV dosage and volumes, normality of data could not be established. Thus, comparisons between those groups were performed using a Brown-Forsythe and Welch ANOVA. Comparisons between scAAV9-GFP and scAAV9-IGF1 were performed using two-way ANOVA with Šidák’s multiple comparison test when interactions were detected. Differences in limb necrosis were analyzed using a Mann-Whitney U test. All statistical analysis was performed in GraphPad Prism (Version 9.0) except for Mann-Whitney U test’s which were performed using Vasserstats.net. In all cases, *P* < 0.05 was considered statistically significant.

## RESULTS

### Optimization of dosage and volume for AAV delivery to the critically ischemic limb

The first goal of this study was to determine the the appropriate AAV dosage and injection volume to effectively transduce the critically ischemic limb without adverse impact on the limb health. To do this, four injection strategies using different injection volumes and viral dosage were compared as shown in Figure 1A. Importantly, AAV’s were delivered to the hindlimb muscle seven days after FAL, a time where the ischemic pathology is significant (24,25,29,31,37). Quantification of the integral of the GFP intensity and area demonstrated a significant effect of viral titer (*P*<0.0001), with groups that received a higher AAV dosage (1.12E+12 viral genomes) having greater GFP infection (Figure 1B,C). There was no significant effect of injection volume on the GFP infection (*P*>0.05). Interestingly, the FDB muscle, located on the pad of the foot, proved to have poor infection regardless of titer or volume delivered. Importantly, neither AAV dosage nor volume altered perfusion recovery in the paw or gastrocnemius muscle when compared to mice that received sham injections (insertion of the needle only) (Figure 1D).

### AAV injection dose dose not adversely impact myofiber size, histopathology, or contractile function in critically ischemic muscle

An assessment of safety is a crucial step for the development of new gene therapies in PAD/CLTI. As such, we investigated whether injection volume of dosing impacted the muscle histopathology or function. Representative images of the limb muscles harvested at the time of euthansia (35 days post-FAL) are shown in Figure 2A. Compared to sham injected muscles, scAAV-GFP delivery did not impact the mean myofiber CSA (Figure 2B). Myofiber size distributions, shown in Figure 2C, demonstrate a greater proportion of smaller myofiber sizes compared to non-ischemic control muscle. Similar to the mean myofiber CSA, myofiber distributions in AAV-treated muscles were consistent with sham injected muscles. Furthermore, quantification of the ischemic lesion area (amount of tissue with evidence of injury) and non-myofiber areas were unaffected by AAV delivery (all *P*>0.05, Figure 2D,E). Because muscle function is an important contributor to PAD/CLTI pathophysiology, we also assessed muscle contractile function using nerve-mediated electrical stimulation of the tibialis anterior muscle. Compared to sham injected mice, there were no significant differences in the maximal absolute (Figure 3A) of specific (Figure 3B) twitch or tetanic forces in mice that received scAAV-GFP. Similarly, no differences in hindlimb grip strength were detected (Figure 3C).

**Figure 2.**
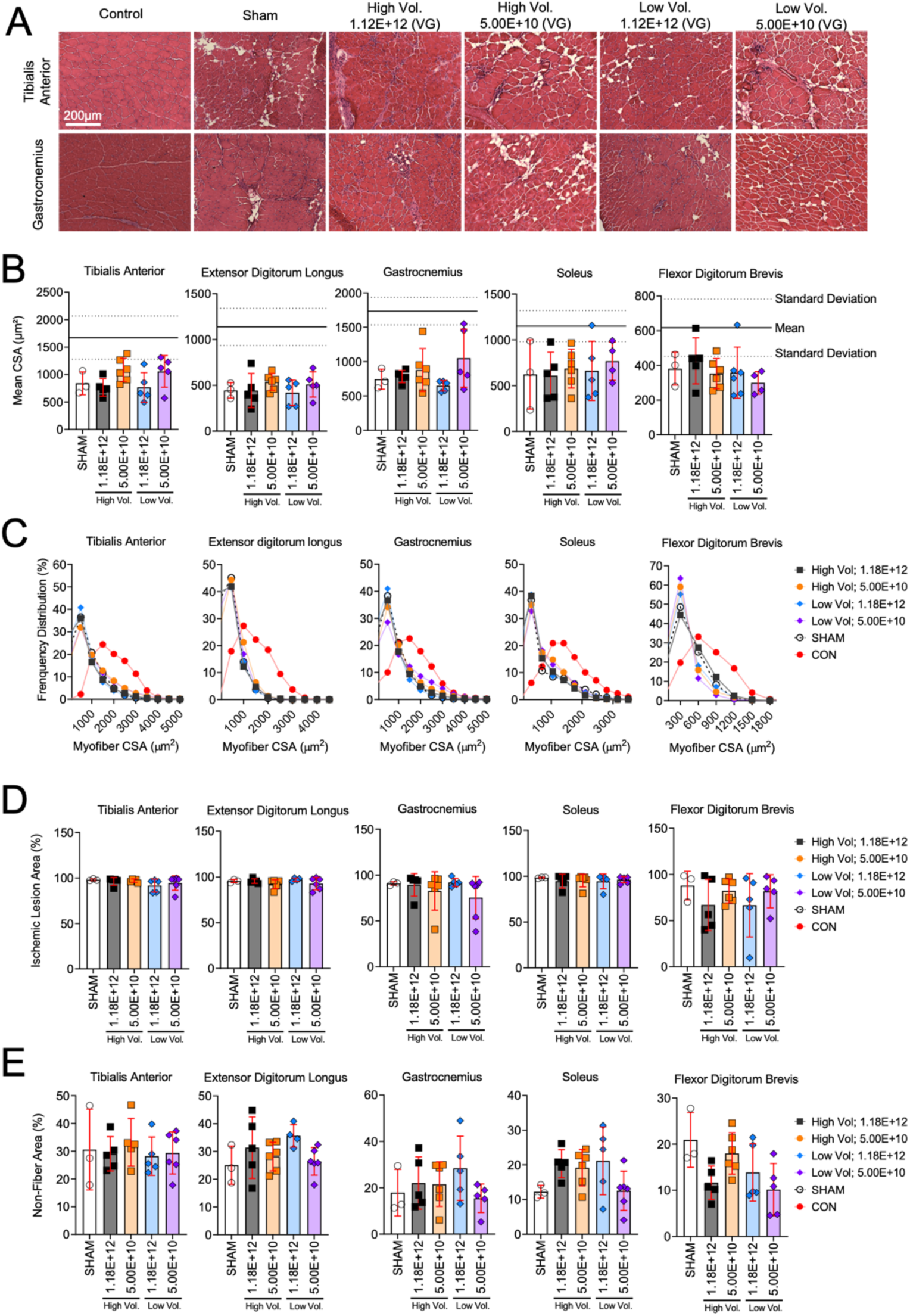
AAV injection does not alter myofiber size or histopathology in critically ischemic muscle. (A) Representative H&E images of tibialis anterior and gastrocnemius muscle injected with scAAV9-CMV-GFP. (B) Quantification of mean of myofiber area of hindlimb muscles (n=3-6/group). (C) Histograms showing distributions of myofiber in hindlimb muscles. (D) Quantification of the ischemic lesion area (percentage of muscle injured). (E) Quantification of non-fiber area (percentage of total muscle area). Error bars represent SD. Panels B, D, and E were analyzed using Brown-Forsythe and Welch ANOVA. The solid line represents the mean and dotted lines the SD of non-ischemic muscle.

**Figure 3.**
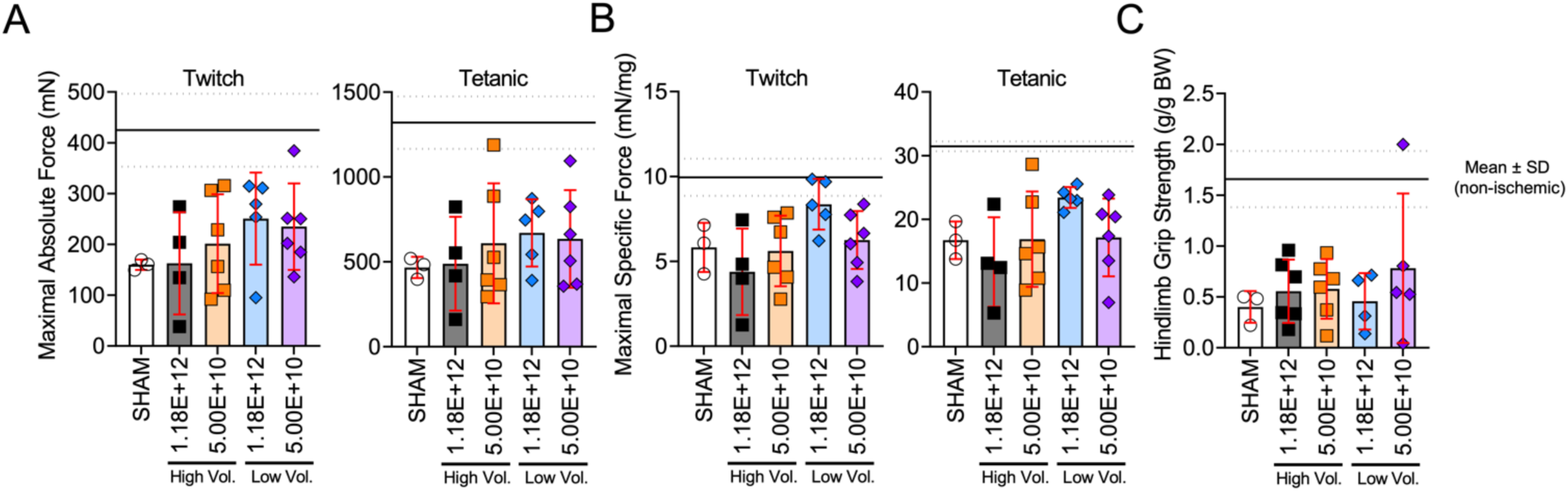
Muscle function was unaffected delivery of AAV to critically ischemic muscle. (A) Maximal absolute twitch and tetanic force of the ischemic tibialis anterior muscle stimulated via the sciatic nerve (n=3-6/group). (B) Maximal specific twitch and tetanic force of the ischemic tibialis anterior muscle stimulated via the sciatic nerve (n=3-6/group). (C) Hindlimb grip strength (normalized to body weight) measured 35 days after surgery (n=3-6/group). The solid line represents the mean and dotted lines the SD of non-ischemic muscle. Error bars represent SD. Panels were analyzed using Brown-Forsythe and Welch ANOVA.

### Treatment with scAAV9-IGF1 dose not improve perfusion recovery or capillary density in the critically ischemic limb

Having established a strategy for delivery AAV-based therapies to the critically ischemic murine limb that allows robust expression without adverse impact, the next step was to investigate the potential efficacy of a therapy. To this end, we chose to determine if treatment of the critically ischemic limb with IGF1 could improve ischemic limb pathology. IGF1 was chosen because of its well-documented ability to promote muscle hypertrophy, enhance muscle regeneration after injury (42,43), and induce angiogenesis (44,45). Figure 4A illustrates the study design employed to determine the therapeutic efficacy of scAAV9-IGF1 treatment delivered to the critically ischemic limb. Analysis of IGF1 mRNA levels in the ischemic limb muscle levels confirmed a signficant increase expression (Figure 4B). Interestingly, depsite performing intramuscular injections only on the surgical limb, the contralateral control limb muscle also displayed an increase in IGF1 mRNA levels. This finding indicates that, despite low levels of limb perfusion at the time of injection, some AAV delivered intramuscularly can enter the circulation and reach tissues distant from the injection site.

**Figure 4.**
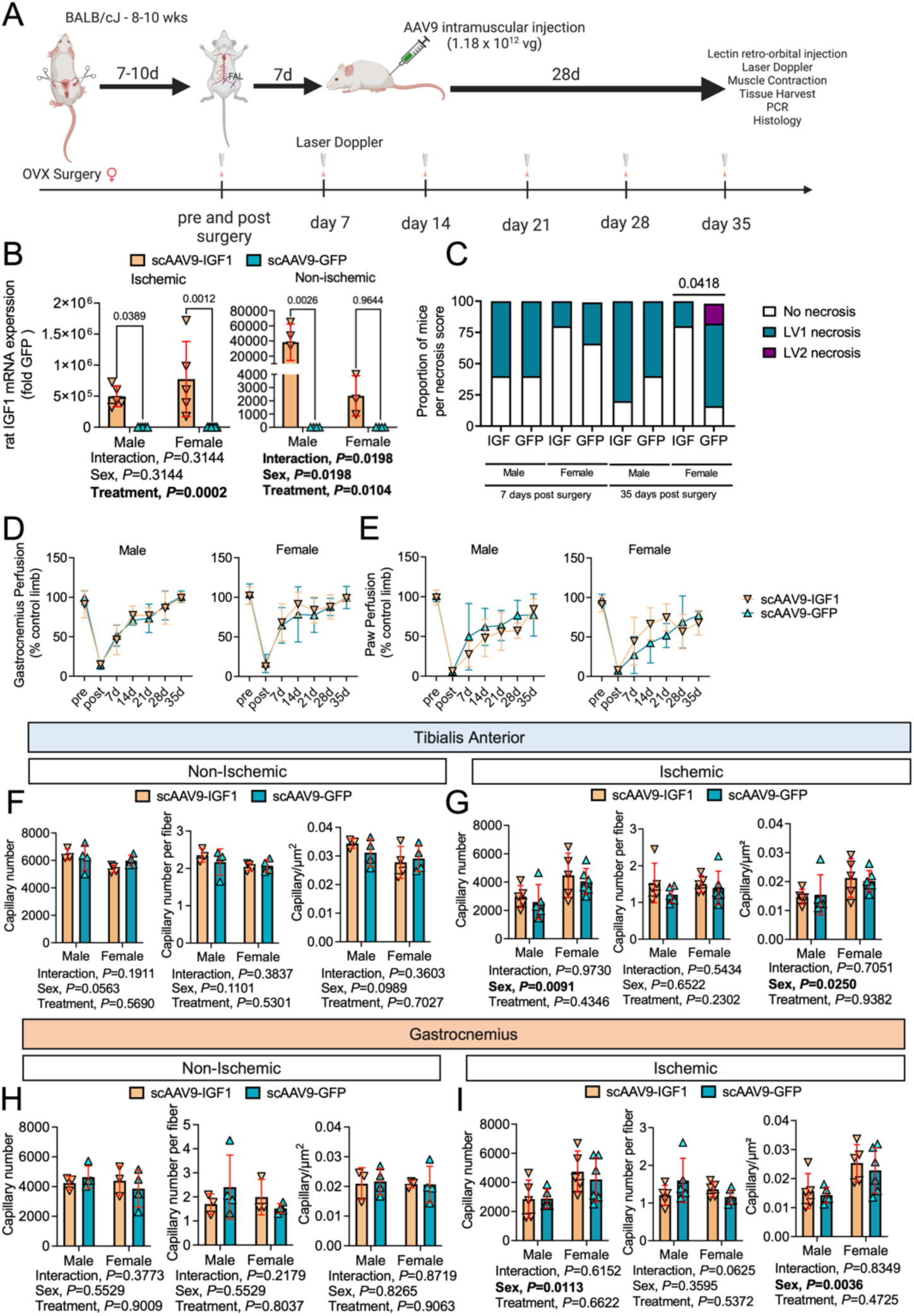
Treatment with scAAV9-IGF1 does not improve perfusion recovery or capillary density in the critically ischemic limb. (A) Graphical abstract of study design created using Biorender.com.(B) Distribution of limb necrosis scores at 7 and 35-days post-FAL surgery (n=5-6/group/sex). (C) mRNA levels of rat IGF1 in the ischemic and non-ischemic gastrocnemius muscle (n=3-6/group/sex). (D) Gastrocnemius perfusion recovery and (E) paw perfusion recovery (n=5-6/group/sex). Quantification of perfused capillary number, perfused capillaries per myofiber, and perfused capillary number per muscle area (μm^2^) within the (F) non-ischemic and (G) ischemic tibialis anterior muscle (n=3-6/group/sex). Quantification of perfused capillary number, perfused capillaries per myofiber, and perfused capillary number per muscle area (μm^2^) within the (H) non-ischemic and (I) ischemic gastrocnemius muscle (n=3-6/group/sex). Error bars represent SD. Panel B was analyzed using Mann-Whitney U test, panels D-I were analyzed using two-way ANOVA.

Female mice that received scAAV9-IGF1 treatment had significantly lower necrosis scores at the time of euthanasia compared to scAAV9-GFP treated mice, indicating that IGF1 therapy may attenuate the progression of limb necrosis (Figure 4C). Laser Doppler measurements found no differences in the perfusion recovery of the gastrocnemius (Figure 4D) or paw (Figure 4E) between groups for both sexes. Consistent with this finding, perfused capillary density was not impacted by the treatment in either the control or ischemic tibialis anterior (Figure 4F,G) and gastrocnemius (Figure 4H,I) muscles. Notably, there were signficant sex effects detected for the perfused capillary density in the ischemic muscles indicating a more robust neovascularization response in female mice when compared to male mice, regardless of treatment group (Figure 4G and 4I).

### Treatment with scAAV9-IGF1 significantly increased muscle size and force production in the critically ischemic limb

Mice that received scAAV9-IGF1 treatment had significantly larger (∼36%) muscle mass in the tibialis anterior (*P*=0.015) and gastrocnemius (*P*=0.045) muscles compared to mice that received scAAV9-GFP (Figure 5A). The extensor digitorum longus was 26% and 33% larger in male and female scAAV9-IGF1 treated mice, although the treatment effect was not statistically significant (*P*=0.064). The mass of deeper-lying plantarflexor muscles including the soleus and plantaris, as well as the flexor digitorum brevis were not different between groups. Functionally, there was a significant treatment effect for maximal absolute twitch (*P*=0.003) and tetanic (*P*=0.008) forces, establishing that scAAV-IGF1 improved muscle strength in mice with PAD/CLTI (Figure 5B). Interestingly, a significant interaction (*P*=0.0418) was detected in the maximal specific force (Figure 5C), but the treatment effect was non-significant (*P*=0.089). Post hoc testing revealed that female mice treated with scAAV-IGF1 had a significantly greater specific force levels compared to scAAV9-GFP female mice (*P*=0.016). A similar trend was observation in the maximal specific twitch forces, although the interaction was not statistically significant (*P*=0.068).

**Figure 5.**
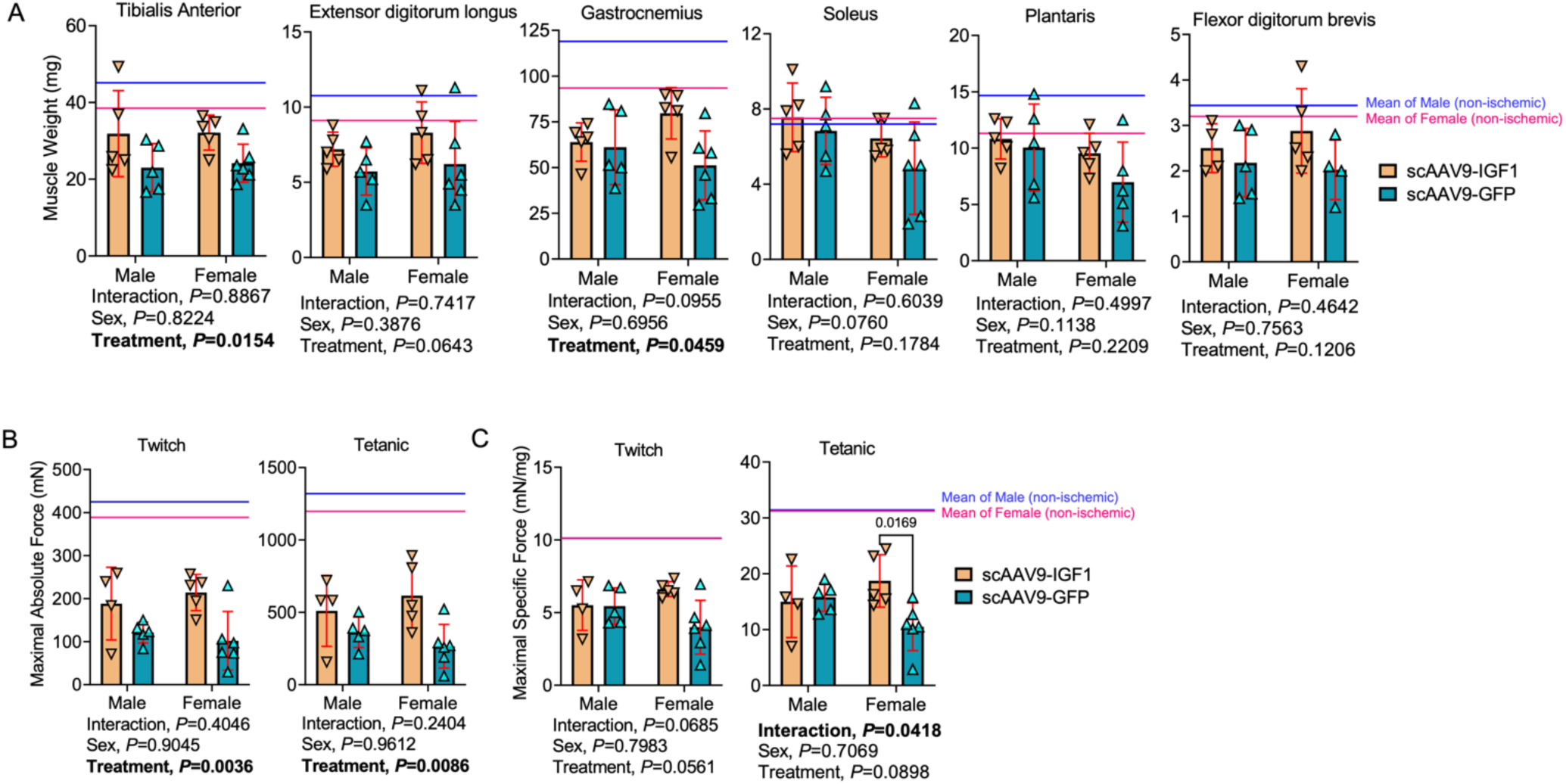
Treatment with scAAV9-IGF1 significantly increases muscle size and force production in the critically ischemic limb. (A) Quantification of muscle weights in mice (n=5-6/group/sex). The solid blue and pink lines represent the mean non-ischemic muscles from male and female mice respectively. (B) Maximal absolute twitch and tetanic force of the ischemic tibialis anterior muscle stimulated via the sciatic nerve (n=5-6/group/sex). (C) Maximal specific twitch and tetanic force of the ischemic tibialis anterior muscle stimulated via the sciatic nerve (n=5-6/group/sex). Error bars represent SD. Panels were analyzed using two-way ANOVA with Šidák’s multiple comparison test when necessary. The solid line represents the mean values for non-ischemic muscle in each sex (blue = male, pink = female).

To access myofiber size and histopathology, immunofluorescence microscopy and hematoxylin and eosin (H&E) staining was performed on transverse sections of the tibialis anterior and gastrocnemius muscles. Frequency distributions of myofiber CSA of the tibialis anterior muscle are shown in Figure 6A. Consistent with muscle weight and absolute force production, a significant treatment effect was observed in the mean myofiber CSA for the tibialis anterior muscle (*P*=0.029, Figure 6B). However, quantification of the non-myofiber area, ischemic lesion area, and the percentage of fibers with centralized nuclei in the tibialis anterior muscle at day 35 post-FAL were not different between groups (Figure 6C-E). Representative images of the tibialis anterior muscle from each group are shown in Figure 6F. Despite an increase in muscle weight, no treatment effect was detected in the ischemic gastrocnemius mean myofiber CSA (*P*=0.31) Figure 6G,H). However, a significant interaction was detected for the mean myofiber CSA (*P*=0.047), and subsequent post hoc testing revealed a non-significant increase in myofiber CSA in female mice treated scAAV-IGF1 (*P*=0.068). Similar interactions, but not treatment effects, were detected for the non-myofiber area (Figure 6I), ischemic lesion area (Figure 6J), and centralized nuclei (Figure 6K) in the gastrocnemius muscle. Post hoc testing again revealed sex-dependent effects of scAAV-IGF1 therapy with females have lower non-myofiber (*P*=0.053), ischemic lesion areas (*P*=0.006), and centrally nucleated myofibers (*P*=0.008) compared to scAAV9-GFP treated females. Representative images of the gastrocnemius muscles from the surgical limb are shown in Figure 6L.

**Figure 6.**
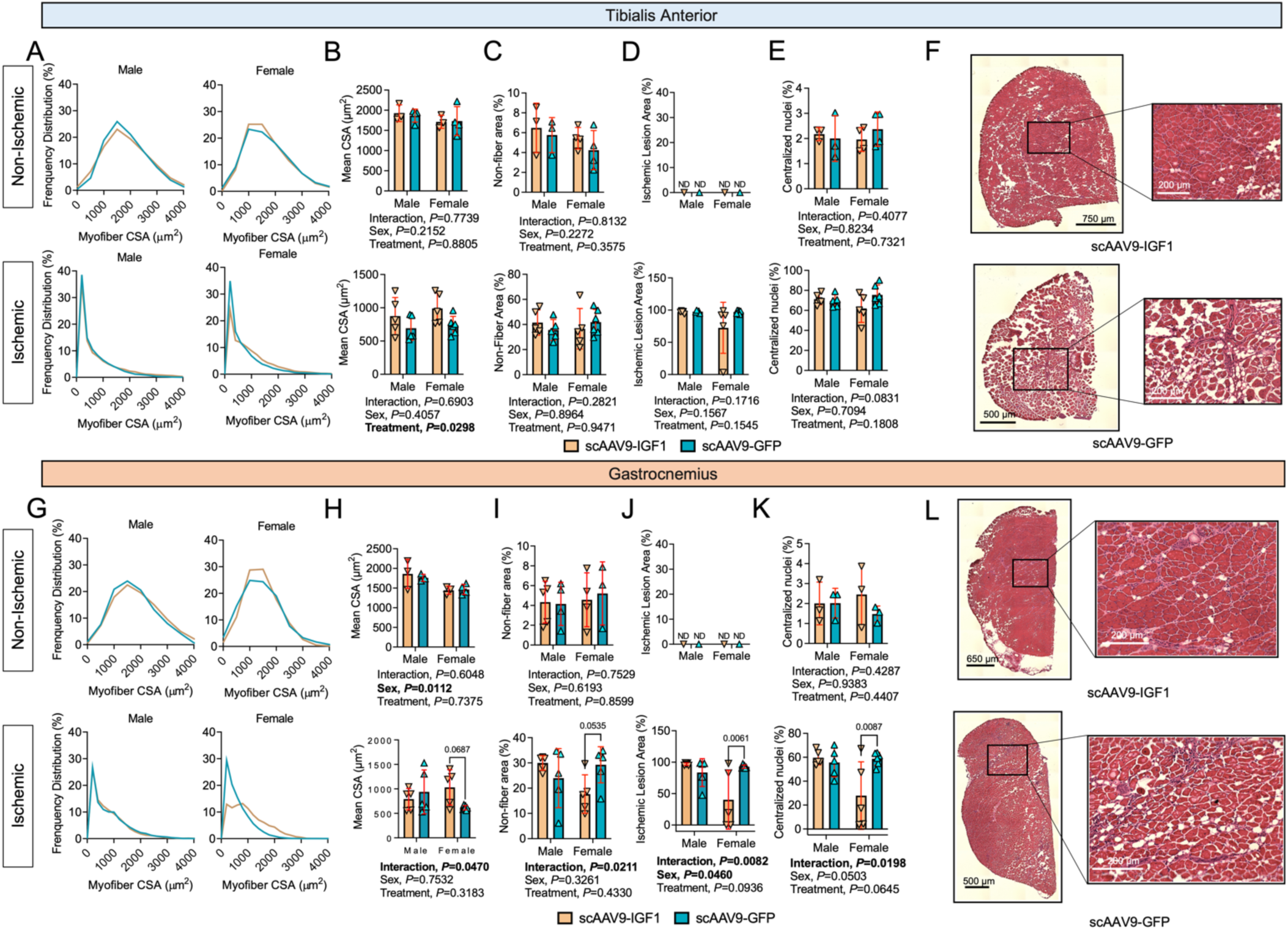
Impact of scAAV9-IGF1 on myofiber size and histopathology in the critically ischemic limb. (A) Histograms of the distribution of myofiber CSA, (B) Quantification of the mean myofiber area, (C) Quantification of non-fiber area (percentage of total muscle area), (D) Quantification of the ischemic lesion (percentage of muscle injured), and (E) Quantification of the percentage of myofibers with centralized nuclei within the tibialis anterior muscles (n=3-6/group/sex). (F) Representative H&E images of tibialis anterior muscles in female mice. (G) Histograms of the distribution of myofiber CSA, (H) Quantification of the mean myofiber area, (I) Quantification of non-fiber area (percentage of total muscle area), (J) Quantification of the ischemic lesion (percentage of muscle injured), and (K) Quantification of the percentage of myofibers with centralized nuclei within the gastrocnemius muscles (n=3-6/group/sex). (L) Representative H&E images of lateral gastrocnemius muscles in female mice. Error bars represent SD. Panels B-E and H-K were analyzed using two-way ANOVA with Šidák’s multiple comparison test when necessary.

## DISCUSSION

Compared to other acquired cardiovascular diseases, therapeutic development for the treatment of PAD/CLTI has been less than desirable. Medical management of PAD routinely involves treatments that reduce the risk of a cardiovascular events, but the only medication approved specifically for the treatment PAD/CLTI is cilostazol which was approved for use in 1999. In this study, we first established an effective and safe approach to deliver AAV-based gene therapy in an experimental model of severe PAD or CLTI. Using the most effective delivery method, the therapeutic efficacy of a gene therapy encoding IGF1, a well-known growth factor that enhances myofiber regeneration (46), was investigated. There are several noteworthy findings from the current study. First, robust infection of the critically ischemic limb can be achieved with scAAV9 without adverse impact on limb pathophysiology. Second, treatment with scAAV9-IGF1 does not improve perfusion recovery or perfused capillary density in the critically ischemic limb. Third, treatment with scAAV9-IGF1 significantly increased muscle size and absolute force production in the critically ischemic limb. Finally, treatment effects with scAAV9-IGF1 displayed some sex-dependent effects with female mice (but not males) having greater specific force and signs of improved histopathology in critically ischemic muscles.

In recent years, gene therapy has gained momentum due to several treatments receiving regulatory approval for the treatment of human diseases. Many gene therapy clinical trials are currently underway or have been recently completed covering a range of diseases including inflammatory, neurological, cancer, infectious, and cardiovascular diseases (47). Numerous cell and gene-based therapies, primary centered in stimulating vascular growth, have been tested in PAD/CLTI patients. Unfortunately, no such therapy has achieved success to date and treatment options remain limited (48). There are numerous factors that have likely contributed to the failed translation of PAD/CLTI therapies from pre-clinical research models to patients. First, PAD/CLTI patients often suffer from multiple comorbidities that accelerate disease pathogenesis and worsen outcomes. The interactions between the patient’s individual genome, environment, and comorbidities are extremely difficult to model in a pre-clinical setting. Second, the primary focus of most PAD/CLTI gene or cell-based therapies has been focused on stimulating blood vessel growth. While this is undoubtedly an important goal, our understanding of the complex interactions between cell types within the PAD/CLTI limb required to achieve effective promotion of both blood vessel growth and regeneration of other injured cells (i.e., muscle, nerve, skin) is incomplete. Moreover, the local microenvironment in the PAD/CLTI limb is quite different from that encountered in other pathologies where gene/cell-based therapies are having success, so PAD/CLTI-specific delivery strategies must be developed.

AAV has emerged as the leading platform for gene therapy in the treatment of human disease. Although generally considered safe and less immunogenic than other viral vectors, there is evidence that wild-type AAV exposure may contribute to production of neutralizing antibodies that may affect gene transfer (49). Additionally, the health of the recipient tissue may also play a role in the efficacy of AAV gene transfer. In a recent study, Mollard *et al*. reported AAV transduction was much lower in muscles regenerating from either cardiotoxin injection of from the effects of muscular dystrophy (50). This effect was observed when AAV treatment was applied both 3- and 45-weeks after cardiotoxin injury. This is a particularly relevant discovery for PAD/CLTI AAV-therapies considering that the ischemic muscle is pathological and shows histological characteristics of regenerating muscle. In the current study, we established efficacious dosing and volumes for the delivery of self-complimentary AAV to the CLTI limb. Robust expression of the GFP transgene was achieved across nearly all hindlimb muscles despite providing the AAV-therapy seven days following FAL, a time where the tissue pathology in BALB/c mice is substantial (24,25,28,29,31,37). Importantly, transgene delivery and expression had no adverse impact on limb perfusion recovery, muscle contractile function, or limb histopathology. In consideration for the need for rapid expression of AAV-therapy in the PAD/CLTI limb, we employed self-complimentary AAV’s to circumvent the need to synthesize the second strand of DNA prior to expression of the transgene.

Although the etiology of PAD/CLTI undoubtedly begins in the vasculature with impaired blood flow to the limb, muscle function/quality has been identified as a major predictor of patient outcomes and quality of life (51-61). Walking performance, as measured by the six-minute walk distance, is widely used to evaluate therapeutic interventions in peripheral artery disease (62-65). A component of walking performance is related to muscular strength, power, and endurance. Various groups have documented pathological abnormalities within the skeletal muscle of PAD patients (66-80). Few clinical studies have employed objective measures of muscle strength or power in PAD patients, particularly in the context of randomized clinical trials. Cross-sectional studies have reported that PAD patients are ∼15-20% weaker than matched adults without PAD (81,82). Further to this, leg strength measures were reported to have strong associations with ABI (82). Two longitudinal observational studies in PAD patients reported that low muscle strength was associated with higher all-cause mortality (60,74). Furthermore, one recent study reported a strong positive association between walking performance and calf muscle myofiber size (83). Taken together, these findings suggest that therapies aimed to increase muscle size/strength may benefit PAD patients. It is difficult to directly compare our results with nerve-mediated contraction in mice to voluntary strength measures in patients. Nonetheless, some descriptive comparisons highlight both the severity of myopathy and magnitude of therapeutic benefit of IGF1 observed herein. Compared to the non-surgical limb, muscle force levels decreased by ∼75% in the FAL limb of scAAV9-GFP mice demonstrating a severe limb myopathy in these CLTI mice. Treatment with scAAV9-IGF1 increase muscle force levels by 40% in males and 230% in females, demonstrating a considerable improvement in muscle function. Considering the magnitude of weakness in PAD patients (∼15-20%), it is reasonable to hypothesize that the achievement of similar increases in muscle strength in human PAD/CLTI patients may significantly improve limb function and quality of life.

Numerous studies have demonstrated that the growth factor IGF-1 can promote muscle protein synthesis resulting muscle hypertrophy (84). IGF-1 has been shown to enhance muscle regeneration from injury or disuse (85-87). In addition to promoting myogenesis, IGF1 has also been proven to induce angiogenesis (45,88), decrease atherosclerosis (89,90), reduce degeneration of motor neurons (91), and suppress inflammation (89,90). In the current study, self-complimentary AAV9-IGF1 therapy significantly increased both muscle size (∼36%) and strength in male and female mice with experimental PAD/CLTI (Figure 5). Interestingly, the therapeutic efficacy of scAAV9-IGF1 was greater in female mice compared to males. For example, male mice treated with scAAV9-IGF1 had ∼40% higher absolute muscle force compared to scAAV9-GFP controls, whereas female mice treated with scAAV9-IGF1 had ∼230% higher muscle force compared to controls. Notably, the observed increase in muscle strength occurred without improvements in perfusion recovery or capillary density (Figure 4). A previous study by Borselli *et al*., (35) reported that treatment with alginate gels contain recombinant VEGF and IGF1 could improve muscle regeneration and force levels after FAL. Similar to results herein, this previous study reported that gels containing only IGF1 did not improve limb perfusion recovery. In contrast to the current study, only mice that received a combination of VEGF and IGF1, but not those that received IGF1 alone, displayed greater muscle size (∼25%) compared to controls. Unfortunately, this previous study only reported muscle force/strength measures for mice treated with VEGF+IGF1, so direct comparisons to the current study regarding the efficacy cannot be made. There are some noteworthy differences in experimental approach between the present study and that by Borselli *et al*. First, the current study used male and female mice, whereas only females were used in the work by Borselli *et al*. Moreover, female mice in the current study underwent oophorectomy to better mimic the post-menopausal condition of female PAD patients. Additionally, the current study utilized BALB/c mice which experience more severe muscle injury, less perfusion recovery, and less angiogenesis following limb ischemia compared to the C57BL/6 mice (31,33). In addition, there are differences both in the delivery method (alginate gel vs. scAAV9) and timing (immediately after FAL vs. seven-days post-FAL) that contribute to a different local microenvironment at the time of treatment.

In the present study, we uncovered some unexpected sex differences regarding the efficacy of the IGF-1 treatment in the critically ischemic limb. As discussed above, the relative improvement in muscle strength in female mice was approximately five times greater than male mice treated with scAAV-IGF1. In the ischemic gastrocnemius muscle, female (but not male) mice treated with scAAV-IGF1 had lower ischemic lesion areas, number of fibers with centralized nuclei, and non-myofiber areas compared with scAAV9-GFP control mice. There are several factors that could explain the sex-dependent efficacy differences. First, estradiol has been shown to rapidly activate the IGF1-receptor (92), as well as stimulate hepatic IGF1 production (93) demonstrating coupling between the estrogen and IGF1 signaling pathways. Female mice in this study had their ovaries removed 7-10 days prior to FAL surgery, which reduces estradiol levels. Further to this, estradiol has also been linked to growth hormone levels (94), which is known to stimulate hepatic IGF1 production. As recently reviewed by McMillin *et al*. (95), disruptions in sex-hormone levels, including deficiencies in estradiol, have been linked to an accelerated loss of skeletal muscle. Thus, the oophorectomy procedure likely makes female mice more vulnerable to ischemic myopathy. This statement is partially supported by our recent study in female BALB/c mice without oophorectomy where the loss of muscle force was ∼50% in the FAL-limb (96), compared to the ∼70% decrease in muscle force reported in the current study with oophorectomized female BALB/c mice. Taken together, these findings could explain the greater efficacy of scAAV9-IGF1 in female mice observed herein. More importantly, the results herein highlight the critical need for adequate female representation in clinical trials enrolling PAD/CLTI patients and the necessity for thorough statistical analysis of sex-dependent treatment efficacies.

### Study Limitations

There are some limitations of the experimental approaches used in this study. First, this study employed younger mice, despite the widely known increase in PAD prevalence with increasing age. Younger mice were used due to the difficulties in obtaining aged mice, especially for strains like BALB/c, where aged colonies are not available from commercial or public vendors. Further to this, PAD/CLTI patients often suffer from numerous co-morbid conditions such as hypertension, diabetes, renal disease, hyperlipidemia that were not present in the mice used in this study. These comorbidities coupled with the genetic and environmental risk factors (i.e., smoking, poor diet, lack of physical activity) are incredibly challenging to replicate in pre-clinical research. Further to this, PAD/CLTI is typically a progressive disease by which atherosclerosis develops slowly over time. In contrast, a rapid and severe loss of limb blood was induced by FAL in the current study.

## CONCLUSIONS

Herein, we aimed to establish a robust and effective method for delivering AAV-based therapies to the critically ischemic limb safely. We provide an in-depth validation of viral infection and assess its safety in mice with experimental PAD/CLTI. Further, we establish the potent therapeutic efficacy of scAAV9-IGF1 in an experimental model of PAD/CLTI which improved both muscle size and strength. Finally, we unexpectedly uncovered sex differences in the effectiveness of IGF1 therapy with female mice exhibiting more powerful improvements in limb function compared to their male counterparts.

## CLINICAL PERSPECTIVES

### Competency in Medical Knowledge

Although skeletal muscle pathophysiology has emerged as a key factor in PAD outcomes, current medical therapies are not directed toward improving limb function. In this study, we establish a safe and effective strategy for delivering AAV-based therapies to the limb muscle in a rodent model of severe PAD/CLTI. Treatment with self-complimentary AAV9-IGF1 was found to significantly increase muscle size and strength in PAD/CLTI mice.

### Translational Outlook

This study demonstrates that therapies aimed to rescue skeletal myopathy in PAD/CLTI can significantly improve limb function and enhance the regenerative ability. Unexpectedly, female mice were found to respond to IGF1 therapy more favorably than male mice. These findings suggest that robust and well-controlled studies in human PAD/CLTI should explore the therapeutic potential of IGF1 therapy, but careful consideration of biological sex is necessary.

## ABBREVIATIONS

AAV =: adeno-associated virus
CLTI =: chronic limb threatening ischemia
CSA =: cross-sectional area
FAL =: femoral artery ligation
GFP =: green fluorescent protein
IGF1 =: insulin-like growth factor 1

